# Calibrating Bayesian decoders of neural spiking activity

**DOI:** 10.1101/2023.11.14.567028

**Authors:** Ganchao Wei, Zeinab Tajik Mansouri, Xiaojing Wang, Ian H. Stevenson

## Abstract

Accurately decoding external variables from observations of neural activity is a major challenge in systems neuroscience. Bayesian decoders, that provide probabilistic estimates, are some of the most widely used. Here we show how, in many common settings, the probabilistic predictions made by traditional Bayesian decoders are overconfident. That is, the estimates for the decoded stimulus or movement variables are more certain than they should be. We then show how Bayesian decoding with latent variables, taking account of low-dimensional shared variability in the observations, can improve calibration, although additional correction for overconfidence is still needed. We examine: 1) decoding the direction of grating stimuli from spike recordings in primary visual cortex in monkeys, 2) decoding movement direction from recordings in primary motor cortex in monkeys, 3) decoding natural images from multi-region recordings in mice, and 4) decoding position from hippocampal recordings in rats. For each setting we characterize the overconfidence, and we describe a possible method to correct miscalibration post-hoc. Properly calibrated Bayesian decoders may alter theoretical results on probabilistic population coding and lead to brain machine interfaces that more accurately reflect confidence levels when identifying external variables.

**Significance Statement:** Bayesian decoding is a statistical technique for making probabilistic predictions about external stimuli or movements based on recordings of neural activity. These predictions may be useful for robust brain machine interfaces or for understanding perceptual or behavioral confidence. However, the probabilities produced by these models do not always match the observed outcomes. Just as a weather forecast predicting a 50% chance of rain may not accurately correspond to an outcome of rain 50% of the time, Bayesian decoders of neural activity can be miscalibrated as well. Here we identify and measure miscalibration of Bayesian decoders for neural spiking activity in a range of experimental settings. We compare multiple statistical models and demonstrate how overconfidence can be corrected.

## Introduction

Decoding, estimating external variables given observations of neural activity, is a fundamental tool in systems neuroscience for understanding what information is present in specific brain signals and areas (deCharms and Zador, 2000; Kriegeskorte and Douglas, 2019). Decoders have been widely used for studying the representation of movement variables, such as speed, force, or position (Humphrey et al., 1970; Georgopoulos et al., 1986), the representation of visual stimuli (Warland et al., 1997; Quiroga and Panzeri, 2009) and the representation of sounds (Theunissen et al., 2004), touch (Diamond et al., 2008), odors (Uchida et al., 2014), and tastes (Lemon and Katz, 2007). Here we examine Bayesian decoders that estimate the probability of each possible stimulus or movement given neural observations (Sanger, 1996; Zhang et al., 1998; Koyama et al., 2010; Chen, 2013). Bayesian models explicitly represent the uncertainty about external variables, and this uncertainty may be useful for understanding perceptual/behavioral confidence (Vilares and Kording, 2011; Meyniel et al., 2015) or for creating more robust brain machine interfaces (Shanechi et al., 2016). However, Bayesian models are not always well calibrated (Degroot and Fienberg, 1983; Draper, 1995). Here we ask whether the uncertainty estimates for Bayesian decoders are correct.

With Bayesian decoders, the conditional probability of stimulus or movement variables given neural responses is calculated using Bayes theorem (Quiroga and Panzeri, 2009). This posterior is the product of a likelihood that describes the probability of neural activity given external variables (an encoding model) and a prior that accounts for other knowledge about the external variable. This framework is very general and can be used to decode categorical or continuous variables in trial-by-trial designs or with continuous time series using spiking timing features or counts as well as other population neural signals (van Bergen et al., 2015; Lu et al., 2021). One common likelihood model for the counts of spiking activity is based on the Poisson distribution and the assumption that the neural responses are conditionally independent given their tuning to the external variable. However, since neural activity has shared (Arieli et al., 1996; Tsodyks et al., 1999) and non-Poisson variability (Amarasingham et al., 2006; Goris et al., 2014), recent studies have focused on better modeling latent structure and dispersion (Scott and Pillow, 2012). Modeling this shared and non-Poisson variability can improve decoding (Graf et al., 2011; Ghanbari et al., 2019).

In this paper, we compare Bayesian decoders with Poisson versus negative binomial noise models as well as decoders with or without latent variables with the goal of understanding how differences in model structure affect the posterior uncertainty. In well calibrated models, the posterior of the external variables should accurately reflect their true probability. For instance, a 95% credible interval – analogous to the confidence interval in frequentist descriptions – should have a 95% chance of containing the true value. However, miscalibration can occur due to model misspecification – when the data is generated by a process that does not match the model assumptions – or when there is unmodeled uncertainty about the model structure (Draper, 1995). Previous studies suggest that neural variability may be an important dimension of the neural code (Urai et al., 2022), and the uncertainty of neural population codes may determine perceptual/behavioral confidence (Knill and Pouget, 2004). Accurate descriptions of population uncertainty in experimental data may, thus, inform for theoretical understanding. In this study, we illustrate the basic problem of miscalibration through simulations and evaluate calibration for experimental data.

We focus on several experimental settings: trial-by-trial decoding of stimulus movement direction from primary visual cortex (V1) and reach direction from primary motor cortex (M1), trial-by-trial decoding of categorical natural images from multiple brain regions, and time-series decoding of animal position from hippocampal recordings (HC). We find that using negative binomial likelihoods and latent variables both improve calibration. However, even with these improvements, Bayesian decoders are overconfident. To solve this problem, we introduce a post-hoc correction for miscalibration that yields more accurate uncertainty estimates.

## Materials and Methods

Code for the results in this paper is available at https://github.com/ihstevenson/latent_bayesian_decoding

### Data

To assess the calibration of Bayesian decoders we use previously collected, publicly available data from 1) macaque primary motor cortex during a center-out reaching task, 2) macaque primary visual cortex during presentation of drifting or static sine-wave gratings, 3) mouse multi-region recordings during presentation of static natural images, and 4) rat hippocampus during running on a linear track.

Data from primary motor cortex (M1) were previously recorded from the arm area of an adult male macaque monkey during center-out reaches. Reaches were made in a 20 × 20cm workspace while the animal was grasping a two-link manipulandum, and single units were recorded using a 100-electrode Utah array (400mm spacing, 1.5 mm length, manually spike sorted manually - Plexon, Inc). On each trial, we analyzed spike counts during the window 150ms before to 350 ms after the speed reached its half-max. Data and additional descriptions of the surgical procedure, behavioral task, and preprocessing are available in Walker and Kording (2013).

Data from primary visual cortex (V1) were previously recorded and shared in the CRCNS PVC-11 dataset (Kohn and Smith, 2016). Single units were recorded using a 96-channel multielectrode array from an anesthetized adult monkey (monkey 3) during presentations of drifting sine-wave gratings (20 trials for each of 12 directions). On each trial we analyzed spike counts between 200 ms and 1.2 s after stimulus onset. Detailed descriptions of the surgical procedure, stimulus presentation, and preprocessing can be found in Smith and Kohn (2008) and Kelly et al. (2010).

We also examine an additional previously recorded, shared dataset from primary visual cortex where stimuli were presented with multiple contrasts (Berens et al., 2012). Here single units were recorded using custom-built tetrodes from an awake male monkey (macacca mulatta). Static sine-wave gratings were presented with different contrasts. Here we use data from subject “D” recorded 2002-04-17. Detailed descriptions of the surgical procedure, stimulus presentation, and preprocessing can be found in Ecker et al. (2010) and Berens et al. (2012).

Multi-region data (ABI) were analyzed from the Allen Institute for Brain Science - Visual Coding Neuropixels dataset (https://portal.brain-map.org/explore/circuits). Detailed descriptions of the surgical procedure, stimulus presentation, and preprocessing can be found in Siegle et al. (2021). Briefly, during the recordings, head-fixed mice were presented with visual stimuli (including Gabor patches, full-field drifting gratings, moving dots, and natural images and movies) while they were free to run on a wheel. We analyze single unit data with spikes sorted from six Neuropixels arrays using Kilosort 2 (electrophysiology session 742951821). Using n=267 single units (742951821, with SNR>3, rate>1 spike/trial) responding to 118 natural images (4873 trials in total).

Data from hippocampus were previously recorded from the dorsal hippocampus of a Long Evans rat and shared in CRCNS hc-3 (Mizuseki et al., 2013). Recordings were made using an 8-shank silicon probe, each shank with 8 recording sites, while the animal ran on a linear track, and single units were automatically spike sorted with KlustaKwik and refined with Klusters. Data from ec014_468 were analyzed in 200 ms bins. Data and additional descriptions of the surgical procedure, behavioral task, and preprocessing are available in Mizuseki et al. (2014)

### Encoding Models

Our goal is to decode an external stimulus or movement variable *x*^∗^ based on spikes observations from *N* neurons 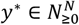 . Here we construct a Bayesian decoder by first fitting an encoding model with training dataset {*x, Y*} where *x* = (*x*_1_…, *x*_*K*_)^’^ denotes the external variable across *K* trials and *y*_*ki*_ (entries of *Y* ∈ *N*^*K*×*N*^) is the number of spikes emitted by neuron *i* during external variable *x*_*k*_. This encoding model allows us to calculate the likelihood distribution *P*(*y*^∗^|*x*^∗^, *x, Y*), and we then use Bayes’ rule to evaluate the posterior distribution *P*(*x*^∗^|*y*^∗^, *x, Y*). In traditional Bayesian decoders, based on generalized linear models (GLMs), the spikes of each neuron are assumed to be conditionally independent given the external variable. Here we examine GLMs with observation models that assume either Poisson noise or negative binomial noise. Additionally, we fit decoders based on generalized linear latent variable models (GLLVMs) where we use the same representation for external variables, but assume the observations are also related or influenced by low-dimensional unobserved variables (i.e., latent variables). GLMs and GLLVMs have been widely used in statistics for modeling count data (McCullagh and Nelder, 1989; Skrondal and Rabe-Hesketh, 2004) and in neuroscience specifically (Brillinger, 1988; Scott and Pillow, 2012).

### Poisson and Negative Binomial GLMs and GLLVMs

The Poisson GLM and negative binomial GLM model the spiking of neuron *i* on trial *k* as *y*_*ki*_ ∼ *Poisson*(*μ*_*ki*_) or *y*_*ki*_ ∼ *NB*(*μ*_*ki*_, *α*_*i*_), respectively, where *Poisson*(*μ*) indicates the Poisson distribution with the rate parameter *μ* and *NB*(*μ, α*) denotes the negative binomial distribution with mean *μ* and variance *μ* + *αμ*^2^. The mean parameter *μ*_*ki*_ in both models is regressed as 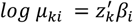 where *z*_*k*_ = *f*(*x*_*k*_) ∈ *R*^*p*^ is a function (e.g. basis expansion) of the external variable *x*_*k*_. For the M1 and V1 decoders we use a Fourier basis to capture the tuning over the circular variable (stimulus or movement direction) *z* = [1 cos *x* sin *x* cos 2*x* sin 2*x*]. For the ABI decoder we simply fit a unique mean for each individual image of the *N* natural image stimuli *z* = [1 1_1_(*x*) ⋯ 1_*N*_(*x*)] where 1_*i*_ (*x*) denotes an indicator function returning 1 when *i* = *x* and 0 otherwise. We estimate *β* and *α* by maximum likelihood estimation (MLE) or, in most cases, maximum a posteriori (MAP) estimation, where we put a Gaussian prior *β*_*j* >1_ ∼ *N*(0, *ηI*) to prevent overfitting (excepting the intercept term). This prior is equivalent to L_2_ regularization.

Since the responses of different neurons may be correlated, the GLM does not generally capture noise correlations - dependencies between neurons beyond what the external variable induces. The GLLVMs extend the GLMs described above by including low dimensional latent factors in the model for the mean parameters. In other words, the Poisson GLLVM and NB GLLVM assume *y*_*ki*_ ∼ *Poisson*(*μ*_*ki*_) or *y*_*ki*_ ∼ *NB*(*μ*_*ki*_, *α*_*i*_) with 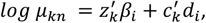, where *c*_*k*_ ∈ *R*^*q*^ is the latent factor for trial *k* (with *q* ≪ *N*) and *d*_*i*_ is the factor loading that describes how the latent states influence neuron *i*.

In this basic form, the latent variable model is not identifiable, and we put several constraints on 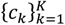 and 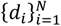 to ensure identifiability. Denote *C* = (*c*_1_, …, *c*_*K*_)^’^ and *D* = (*d*_1_, …, *d*_*N*_), and write the singular value decomposition of *CD* as *CD* = *UΣV*^’^. Following Miller and Carter (2020), we constrain: 1) *U* and *V* to be orthogonal, 2) *Σ* to be diagonal matrix, with diagonal elements > 0 and sorted in descending order and 3) the first nonzero entry for each column of *U* to be positive. Then we let *C* = *UΣ* and *D* = *V*^′^, or equivalently let *C* = *U* and *D* = *ΣV*^′^ . The model parameters then are estimated by maximizing the likelihood via alternating coordinate descent algorithm, i.e. updating the “neuron” part (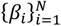 and *D*) and the “latent” part (*C*) until convergence is achieved.

In cases where the number of trials is relatively small, when *p* is large, or when the spiking is extremely sparse, both the GLM and GLLVM can overfit or fail to converge (Zhao and Iyengar, 2010). In addition to the Gaussian prior (i.e. L2 penalty) on *β* we also include a Gaussian prior on *C*, and find the maximum a posteriori (MAP) estimates rather than the MLE.

### Approximate Bayesian Decoding

Once the encoding model is fitted with training data x and y, we then decode the external variable *x*^∗^ based on new observations of spikes *y*^∗^ ∈ *N*^*N*^, by evaluating the posterior distribution *P*(*x*^∗^|*y*^∗^, *x, Y*). For the GLM, we have

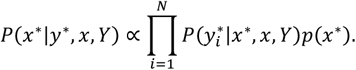

The results here all assume a flat/uniform prior on *p*(*x*^∗^); however, in general, this term can incorporate prior information about the external variables.

For the GLLVM we additionally need to account for the latent variables. Since the data used for fitting the encoding model is not the same as decoding dataset, the latent state *c*_*k*_, depending on specific trials, acts as a nuisance parameter. We then obtain the posterior

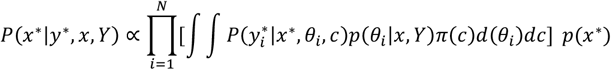

Where *θ* denotes the parameters {*α, β, d*} . When the training set size *K* is small, the parameter estimates for the encoding model can have substantial parameter uncertainty (Cronin et al., 2010). However, in practice, including parameter uncertainty (via MCMC) does not typically affect the posterior over the external variable (see results in Wei, 2023). We thus approximate the full posterior by plugging in the MLE/MAP estimates 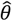.

Our goal is then to calculate the marginal predictive likelihood 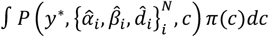. If we assume the observations *y*^∗^ to be conditionally independent given both stimuli and latent factors this is given by 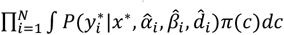. Although there is no closed form solution to the integral, we can use the Laplace approximation, such that

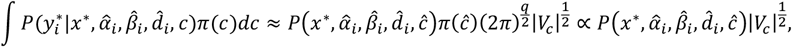

where *ĉ* is the ML (or MAP) estimate and 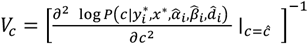

Since the posterior distribution of *x*^∗^ is not necessarily unimodal, we evaluate the posterior distribution by grid approximation, which works efficiently for a one-dimensional case. In other words, we first compute the un-normalized posterior density at a grid of values that cover effective rage of *x*^∗^, and then normalize the density.

### Coverage and Constant Correction

To assess the calibration of these decoders for continuous variables we compare the frequentist coverage (fraction of trials on which the true stimulus/movement falls within a highest density region) to the nominal/desired probability. For a well-calibrated Bayesian model, the highest posterior density (HPD) regions of a given size (e.g. the 95% region) should contain the true values with the nominated probability (e.g. 95%). Here we compute the (cross-validated) proportion of trials for which the true stimulus/movement falls within the HPD regions (the “coverage”) as we vary the size of the credible set.

For categorical posteriors, there are several scoring rules that have been previously described, such as the Brier score (Gneiting and Raftery, 2007), but, here, to emphasize “coverage”, we extend our calculations with continuous credible regions to use discrete credible sets. We construct the HP set, as before, adding the highest probability categories until the probability *m* in the set meets the nominated probability *m*^∗^ with *m* ≥ *m*^∗^. For continuous distributions, credible regions can be calculated so that there are minimal errors between the desired probability (*m*^∗^) and the probability in the credible set (*m*), but for categorical distributions, there can be a substantial mismatch between these quantities. For instance, suppose we want to find the coverage of a 25% credible set, but category 1 has posterior probability 50% on average across trials. To correct for this mismatch, we adjust the empirical coverage for categorical posteriors (ABI results below) by a factor of *m*^∗^/⟨*m*⟩ (e.g., .25/.5 for the example above), where ⟨⋅⟩ denotes an average across trials.

Since most Bayesian decoders appear to be badly calibrated, we consider a post-hoc correction (i.e. recalibration). This correction is similar to the “inflation factor” in ensemble probabilistic forecasting (Wilks, 2002; Gneiting and Raftery, 2007) where similar types of overconfidence can occur (Raftery et al., 2005). Namely, here we consider decoding with a modified posterior *Q*(*x*^∗^|*y*^∗^, *x, Y*) ∝ exp(*h* log *P*(*x*^∗^|*y*^∗^, *x, Y*)) for some constant *h* > 0. Decoding from the modified posterior *Q*(*x*^∗^|*y*^∗^, *x, Y*) does not change the accuracy, but allows the confidence to be adjusted. Here we fit *h* by minimizing the squared error between the empirical and nominal coverage probability over the full range (0, 1).

### Dynamic Models

The GLM and GLLVM described above assume that trials are independent. However, in many cases, it is more appropriate or desirable to decode with a dynamic model. Rather than decoding the external variable on trial *k*, we wish to decode the external variable *x*_*t*_ at time *t* and to incorporate smoothness assumptions relating *x*_*t*_ to previous time points. Such state space models have been previously described for Poisson observations (Smith and Brown, 2003; Paninski et al., 2010; Vidne et al., 2012), and applied for decoding (Lawhern et al., 2010). Here we describe decoding with a dynamic NB GLLVM, for which the Poisson model is a special case (see Wei (2023) for additional detail).

Briefly, we assume that the observation for neuron *i* at time *t* follows

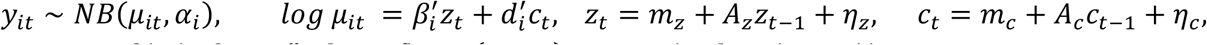

where *z*_*t*_ = *f*(*x*_*t*_), *β*_*i*_ ∈ *R*^*p*^, *d*_*i*_ ∈ *R*^*q*^ and (*η*_*z*_, *η*_*c*_) ∼ *N*_*p*+*q*_(0, *diag*(*Q*_*z*_, *Q*_*c*_)). With initial conditions given by *z*_1_ ∼ *N*(*z*_0_, *Q*_*z*0_) and *c*_1_ ∼ *N*(*c*_0_, *Q*_*c*0_) . To make the model identifiable, we put the same set of constraints on the model parameters as above. Denote *C* = (*c*_1_, …, *c*_*T*_)^’^ and *D* = (*d*_1_, …, *d*_*N*_)^’^, let 1) *C*′*C* be diagonal, with diagonal elements sorted in the descending order, 2) *D*^’^*D* = *I*_*p*_ and 3) the first non-zero entry for each column of *C* is positive.

When fitting the encoding model, {*z*_*t*_ } is observed and {*z*_0_, *Q*_*z*0_, *m*_*z*_, *A*_*z*_, *Q*_*z*_ } do not need to be estimated. We fit the remaining model parameters by a cyclic coordinate descent algorithm, i.e., alternatively updating the “neuron” part 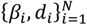 and “latent” part 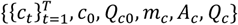. The “latent” part is fitted via an expectation maximization (EM) algorithm with a normal approximation in the E-step, following (Lawhern et al., 2010). For decoding, we plug in the fitted 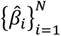 and 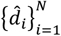 and refit 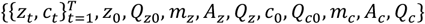 via an EM algorithm again using a normal approximation at E-step. Note that here, 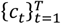 are not treated as nuisance parameters. For the results decoding position from hippocampal activity, we assume that *m*_*z*_ = 0, *m*_*c*_ = 0, *A*_*z*_ = *I*, and *A*_*c*_ = *I*. Additionally, rather than a direct grid approximation for the posterior over *x*^∗^, the posterior is approximated as a multivariate normal distribution over 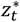 . To assess accuracy and coverage, we evaluate the multivariate normal distribution along a grid in *x*^∗^ for each *t* separately and normalize, 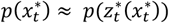.

## Results

Bayesian decoders are based on first fitting tuning curves for each neuron using training data. The encoding model determines the likelihood distribution, and, for traditional (naïve) Bayesian models, neurons are assumed to be conditionally independent given the external variables. During decoding we then use Bayes’ rule to calculate the posterior distribution over possible stimuli or movements given the observed neural activity. Here we focus on assessing not just the decoding accuracy but the uncertainty of the posterior under different models and experimental settings. Our goal is to determine to what extent the traditional models, as well as more recently developed latent variable models, have well-calibrated posterior estimates (i.e., where the posterior probabilities match the true probabilities of the external variable taking specific values).

To illustrate the problem of model calibration we consider a hypothetical set of Bayesian decoders (Fig 1A). The average error is the same for each of these decoders, since the maximum and means of the posteriors are identical, but the uncertainty of the decoders varies. There is underconfidence or overconfidence on single trials, and, across trials, the posterior distributions do not necessarily match the distribution of errors. When errors occur an overconfident decoder will not have proper coverage of the true value. On the other hand, an underconfident decoder will cover the true value too often for the desired confidence level. In our example case, imagining 5 trials and an 80% credible interval, a well-calibrated decoder correctly covers the true value for 4 of 5 trials, while the overconfident decoder only covers 1 of 5 and the underconfident decoder covers 5 of 5. In general, overconfident decoders will have lower coverage than desired, while underconfident decoders will have higher coverage than desired (Fig 1B).

**Figure 1:**
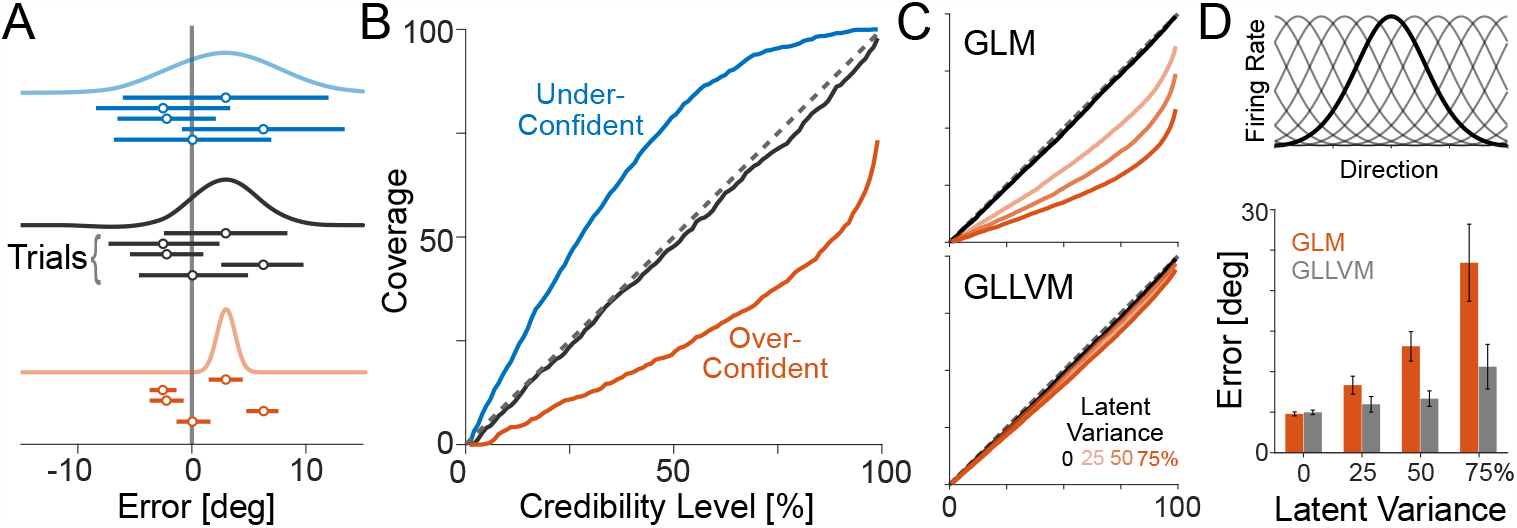
Bayesian decoders can misestimate uncertainty. A) Examples of posteriors for three toy Bayesian decoders: an under-confident (blue), over-confident (red), and a well-calibrated (black) decoder provide posterior estimates for each trial. Curves denote single-trial posteriors and lines below each posterior denote 80% the credible intervals, and credible intervals for an additional four trials. Dots denote MAP estimates. Coverage is measured by whether the highest posterior density regions cover the true value (Error=0, in this case). B) Coverage as a function of the desired confidence level for each decoder. C) In a simulation of homogeneous neurons receiving latent input in addition to their tuning to an external variable, we find that a GLM-based decoder is increasingly over-confident as the contribution of the latent input increases (top). Modeling the latent input with a GLLVM, even though it is unknown, reduces over-confidence (bottom). For clarity, curves are averages of multiple simulations. D) Tuning curves for the simulated population (top) and median cross-validated error for the MAP estimates (bottom) for the GLM (red) and GLLVM (gray) averaged across multiple simulations. Error bars denote standard deviation across simulations.

Bayesian models can have poor calibration when the model is misspecified. To illustrate how such misspecification could occur with neural data we simulate the impact of latent variables on a traditional Bayesian decoder. Here noisy spike observations are generated by a population of identically tuned neurons (Fig 1D, top) with Poisson variability. However, in addition to their stimulus/movement tuning neurons receive a common one-dimensional latent input that increases or decreases activity on individual trials. Since this input is shared by the entire population (of 20 neurons in this case), it produces correlated variability. A traditional Bayesian decoder first fits tuning curves for each individual neuron (here using a Generalized Linear Model - GLM - with Poisson observations). The posterior is calculated assuming that neural responses are conditionally independent given the stimuli, and, as before, we can quantify the coverage by identifying the highest posterior density (HPD) regions. In this more realistic simulation, the posterior can be multimodal resulting in multiple credible regions rather than just a single credible interval. However, since the GLM decoder does not account for the latent variable, the decoder is over-confident (Fig 1C, top) and less accurate (Fig 1D, bottom). When the latent variable has a larger impact on neural responses relative to the impact of the stimulus, errors increase, and the decoder is increasingly overconfident. Hence, traditional Bayesian decoders used in the literature by assuming the independence between responses given the stimuli can have high error and over-confidence in the presence of latent variables.

Modeling the latent variable reduces error and provides well-calibrated posteriors. Here we use a Poisson Generalized Linear Latent Variable Model (GLLVM, see Methods) where the encoding model is fit to account for the tuning curve, as well as the contribution of a shared low dimensional latent variable. Under the GLLVM, neural responses are not conditionally independent given the stimulus. Rather, for each trial the latent variable is estimated, and, during decoding, the latent variable is marginalized over in order to generate the posterior distribution over stimuli. The error for the GLLVM decoder still increases as the latent variable has a larger relative impact on neural responses (Fig 1D, bottom), but the coverage closely follows the desired credibility level (Fig 1C, bottom). Well calibrated decoders (such as the GLLVM in this simulation) have the advantage that the posterior appropriately covers the true stimulus.

To further illustrate how overconfidence arises we consider a single tuned neuron in the GLLVM (Fig 2). Here a neuron is tuned with a preferred stimulus/movement direction of 0 deg. However, a latent variable that changes from trial to trial can shift the tuning curve up or down. This latent variable creates an additional source of ambiguity when a specific spike count is observed. We cannot distinguish between a situation where the neuron is spiking during the presence of a preferred stimulus and a situation where the neuron is spiking during a non-preferred stimulus that coincides with an excitatory latent input. For stimulus *x* and neural responses *y*, the key difference between the GLM and GLLVM decoders is that instead of using the posterior *p*(*x*|*y*) based only on a tuning curve model, we model an additional latent variable *z* and decode from the marginal posterior distribution ∫ *p*(*x*|*y, z*)*p*(*z*)*dz* . Since marginalizing, in general, increases uncertainty, the posterior distributions for individual neurons under the GLLVM will be more uncertain than those of a GLM with the same noise model.

**Figure 2:**
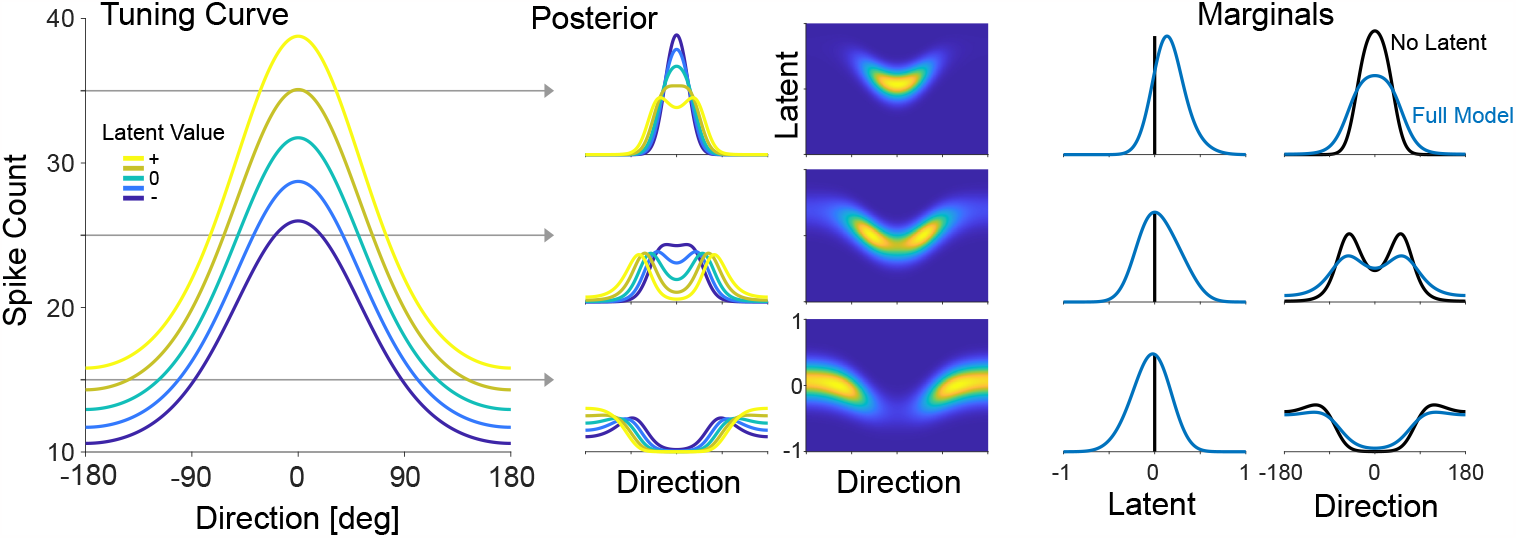
Latent variables increase posterior uncertainty when modeled. A single neuron tuned to reach direction may additionally be impacted by a latent variable (left). After fitting the encoding model, we can find the joint posterior over the value of the latent variable and the reach direction given an observed spike count (middle). To decode the reach direction, we marginalize/integrate over the latent variable (right). The full model (blue) has higher uncertainty for reach direction than a model that does not take the latent variable into account (black).

### Trial-by-Trial Experimental Data

For experimental data we do not know the true model. However, the calibration and accuracy of Bayesian decoders can be assessed empirically. Here we compare GLM and GLLVM Bayesian decoders in three experimental settings: 1) decoding reach direction during a center-out task using recordings from primary motor (M1), 2) decoding sine-wave grating movement direction using recordings from primary visual (V1) cortex, and 3) decoding the identity of a natural image stimulus using multi-region Neuropixels recordings from the Allen Brain Institute (ABI). These data were previously collected and publicly shared (see Methods), and for each setting we evaluate decoding accuracy as well as coverage – the fraction of trials where the true stimulus falls within the highest density regions of the posterior (HPD).

We compare four models 1) Poisson-GLM, 2) negative binomial-GLM, 3) Poisson-GLLVM, and 4) negative binomial GLLVM. For M1 and V1, we model tuning curves using a Fourier basis. For ABI, we model the spike counts in response to each of 118 images and regularize to prevent overfitting (*η* = 100). For the GLLVMs, we model a one-dimensional latent variable that co-modulates the responses of each neuron in the recorded population in addition to the tuning curves. That is, we fit an encoding model which predicts the response of each neuron on each trial as conditionally independent Poisson or negative binomial observations. During decoding we evaluate the posterior distribution over possible external variables and marginalize over the latent variable in the case of the GLLVM. All results are cross-validated (10-fold) such that the decoders are trained on one set of trials and error/accuracy and uncertainty are evaluated on test data.

For experimental data, there is substantial heterogeneity in tuning curves (Fig 3A-C), and posteriors may be continuous or discrete depending on the experimental context. However, as with the toy examples above, the GLLVM (in this case, with a negative binomial observation model) tends to have posteriors with higher uncertainty compared to the GLM (Fig 3E-F). On single trials, the posteriors tend to be wider and to have lower probabilities for the (MAP) point estimate for the GLLVM. In both continuous and discrete cases, outcomes that were assigned near-zero probability under the GLM are assigned non-zero probability under the GLLVM.

**Figure 3:**
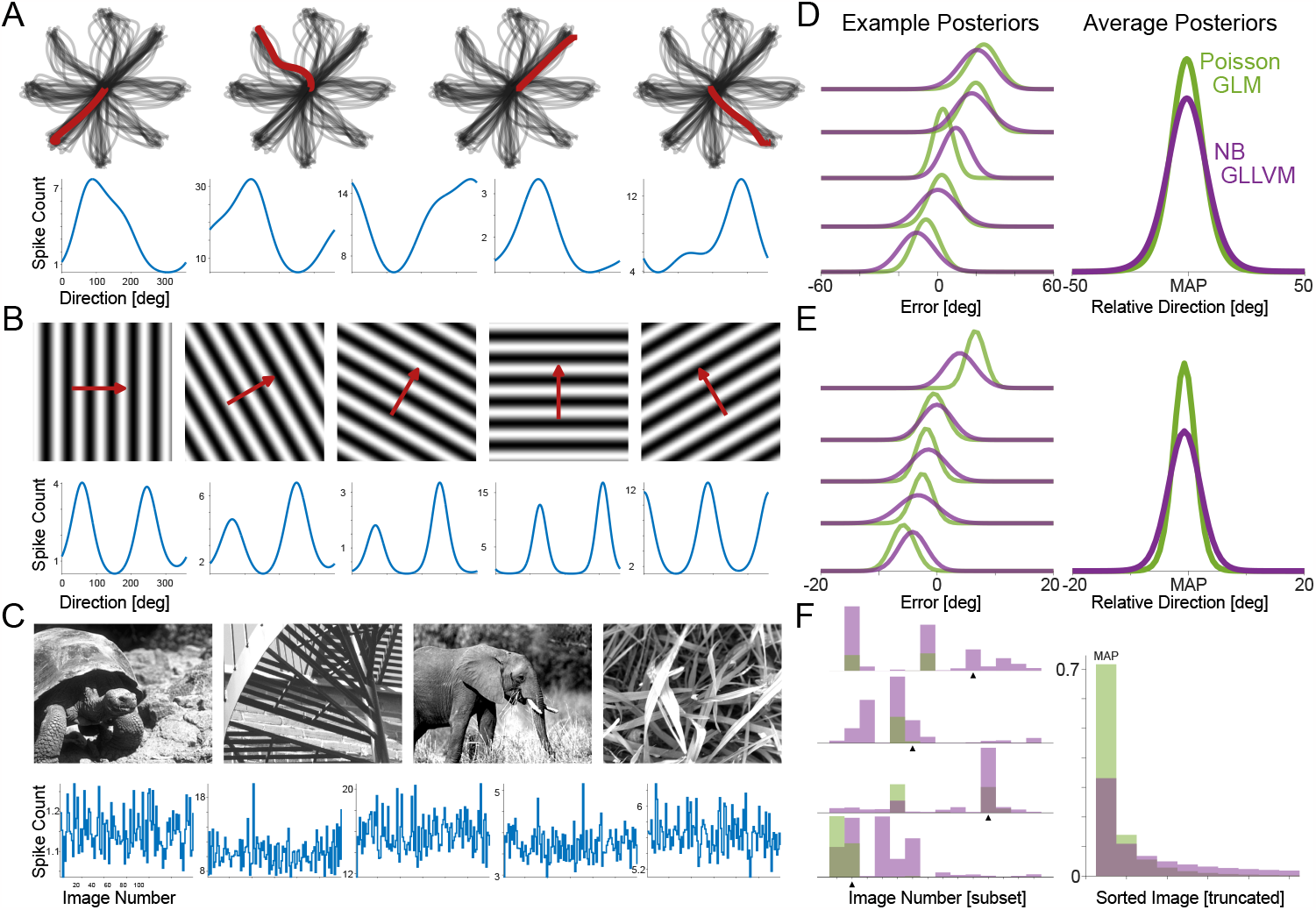
Experimental decoding tasks and example posteriors. A) For M1 data, we aim to decode target direction in single trials of a center-out reaching task, B) For V1 data, we aim to decode stimulus (full-field grating) movement direction in single trials, and C) For ABI data, we aim to decode the identity of a natural image stimulus on single trials. For each case, example stimuli (top) and tuning curves for individual neurons (bottom) from the Poisson GLM fits. (D-F) show example posteriors for single trials (left) as well as the average posterior aligned to the MAP estimate (right). For ABI, note that the posteriors are discrete distributions and, for clarity, only a subset of images are shown. In (F), black triangles denote the true image stimulus.

As with the simulations above, we find that Bayesian decoders tend to be over-confident (Fig 4A-C). For all three experimental settings (M1, V1, and ABI), the highest posterior density (HPD) regions cover the true stimulus/movement less often than desired for all credible levels when decoding from all recorded neurons. For the Poisson GLM, for example, when we specify a 95% credibility level, the posteriors from M1 only include the true target direction 70% of the time, posteriors from V1 only include the true stimulus direction 51%, and posteriors from ABI only include the true natural image stimulus 31% of the time. The negative binomial GLM has better coverage than the Poisson GLM, while adding latent variables improves coverage even more. The best-calibrated model of these four is the negative binomial GLLVM - here when we specify a 95% credibility level, the posteriors from M1 include the true target direction 81% of the time, posteriors from V1 include the true stimulus direction 82%, and posteriors from ABI include the true natural image stimulus 86% of the time. Traditional Bayesian decoders can thus have substantial over-confidence, and calibration is improved by adding latent variables.

**Figure 4:**
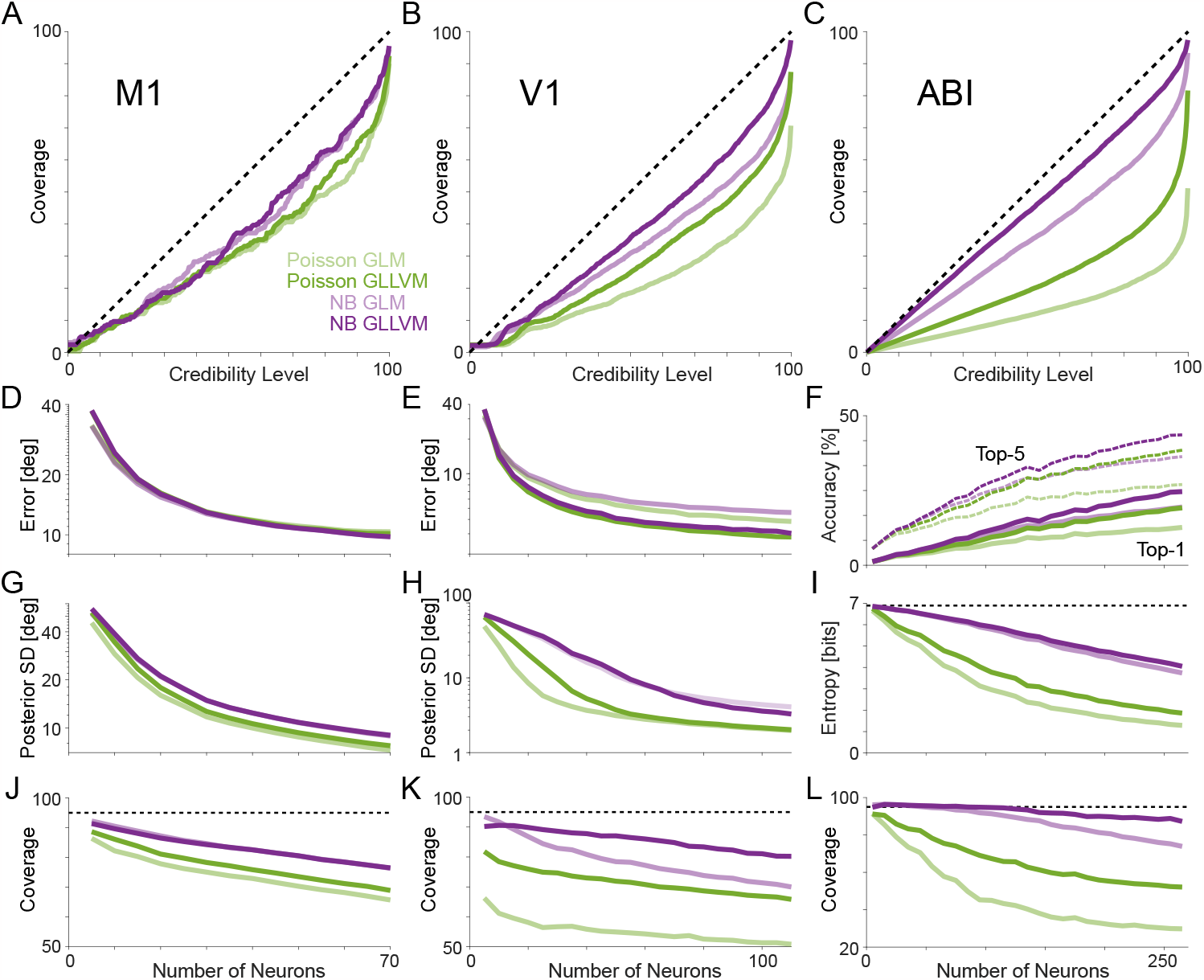
Coverage results for three experimental datasets and four decoders. Decoding reach direction from neurons in M1 during a center-out task (A), decoding stimulus direction from neurons in V1 during presentation of drifting gratings (B) and decoding the identity of a natural image from multiple brain regions (C), Bayesian decoders tend to be over-confident. Latent variable models (P-GLLVM and NB-GLLVM) are better calibrated than their GLM equivalents, and negative binomial models tend to be better calibrated than their Poisson equivalents. Cross-validated error/accuracy (D-F), uncertainty (G-I), and coverage (J-L) each change as a function of how many neurons are included in the model. Accuracy increases with increasing numbers of neurons and uncertainty decreases. However, calibration (the degree of over-confidence) gets worse as more neurons are included in the model. Error in D and E, denotes median error. SD in G and H is circular standard deviation. Dashed line in (I) denotes maximum entropy over the natural images. Dashed lines in J-L denote a nominated 95% coverage. M1 results in D, G, and J are averaged across 200 sets of neurons, V1 results in E, H, and K are averaged across 100 sets of neurons, and ABI results are averaged across 20 sets of neurons.

As previous studies have noted, non-Poisson observation models and latent variables can alter, and in many cases improve, decoding accuracy. Here, for M1 and V1, we calculate the absolute circular distance between the true target/stimulus direction and the maximum a posteriori (MAP) estimate of the target/stimulus direction from the Bayesian decoders on each trial. For ABI, we assess the accuracy based on whether the top-1 or top-5 categories of the discrete posterior include the true stimulus image on each trial. For the full populations of M1 data, the models do not have substantially different errors (median across trials 9.8 deg, 9.5 deg, 9.8 deg and 9.8 deg for the P-GLM, NB-GLM, P-GLLVM, and NB-GLLVM, respectively). For the V1 data, the Poisson GLM outperforms the NB-GLM (median error 3.8 deg vs 4.5 deg, Wilcoxon signed rank test, p<10^-12^, z=7.5), and the Poisson GLLVM outperforms the NB-GLLVM (median error 2.8 deg vs 3.0 deg, Wilcoxon signed rank test p<10^-12^, z=7.7). For ABI data, however, the NB models out-perform the Poisson models (top-1 accuracy 15.6% [14.6, 16.7] for P-GLM vs 23.0% [21.9, 24.2] for NB-GLM). For V1, the GLM-based models have slightly lower error than the GLLVM (p<10^-12^, z=17.0, Wilcoxon signed rank test for Poisson GLM vs GLLVM), but for the ABI data, the GLLVM models improve accuracy substantially (22.3% [22.1, 24.5] for P-GLLVM and 30.1% [29.2, 31.8] for NB-GLLVM). In all cases, for randomly sampled subnetworks, we find that the cross-validated error decreases (or accuracy increases) as a function of how many neurons are included in the decoder for all models (Fig 4D-F).

These error and accuracy measures are based on the MAP estimates of the external variable; however, there are also differences across models in the dispersion of the posteriors. The NB models have higher circular standard deviations than the Poisson models for the M1 and V1 data and substantially higher entropy for ABI (Fig 4G-I). For M1, the circular standard deviation of the posterior is 7.2 deg for the Poisson GLM (median across trials) compared to 8.8 deg for the NB-GLM (p<10^-12^, z=14.3, two-sided Wilcoxon signed rank test), and 7.7 deg and 9.0 deg for the P-GLLVM and NB-GLLVM (p<10^-12^, z=13.9, two-sided Wilcoxon signed rank test). For V1, the median circular standard deviation is 2.0 deg for the P-GLM compared to 4.0 deg for the NB-GLM (p<10^-12^, z=38.8) and 2.0 deg vs 3.3 deg for the P-GLLVM and NB-GLLVM (p<10^-12^, z=-35.0, two-sided Wilcoxon signed rank test). For ABI, the average entropy is 1.26 bits for the P-GLM and 2.7 bits for NB-GLM (t(4872)=136.4, p<10^-12^, paired t-test), 1.8 bits for P-GLLVM, 4.0 bits for NB-GLLVM (t(4872)=19.6, p<10^-12^, paired t-test compared to NB-GLM). In the case of decoding natural images from ABI, the GLLVMs are less certain and more accurate, than the GLMs.

Differences in the dispersions of the posteriors are reflected in differences in coverage. As more neurons are used for decoding the models become increasingly overconfident and badly calibrated (Fig 4J-L), even as the error decreases (Fig 4D-E) or accuracy improves (Fig 4F). The negative binomial GLLVM has the best coverage across datasets and population sizes but note that the coverage is still less than desired (95% for Fig 4J-L).

### Interpreting latent variable models

Including a latent variable allows the GLLVMs to account for variation in neural responses to the same stimulus/movement. Here, with a one-dimensional model, the GLLVM primarily accounts for the overall fluctuations in population activity from trial-to-trial (Fig 5). While the GLM only predicts variation between stimuli/movements for both M1 (Fig 5A) and V1 (Fig 5B), the GLLVM accounts for the fact that some trials tend to have higher overall activity across the population while other trials have lower activity. This trend is apparent when examining the overall population activity – here calculated as the sum of the log activity. We also examine correlations between responses of pairs of neurons (Fig 5, right). Here we calculate stimulus and noise correlations by shuffling responses to the same stimuli/movements. Stimulus correlations reflect the average on shuffled data, while noise correlations are given by the observed correlations minus the shuffled correlations, and, for the models, we sample spike counts to mimic the observed data. Since the GLM assumes that neurons are conditionally independent given the stimulus/movement, it accounts for stimulus correlations but tends to underestimate noise correlations. The GLLVM, on the other hand, accurately accounts for both stimulus and noise correlations. This pattern is present in the overall correlation matrices, as well as, when averaging over pairs of neurons based on the differences in their preferred directions (Δ*PD*).

**Figure 5:**
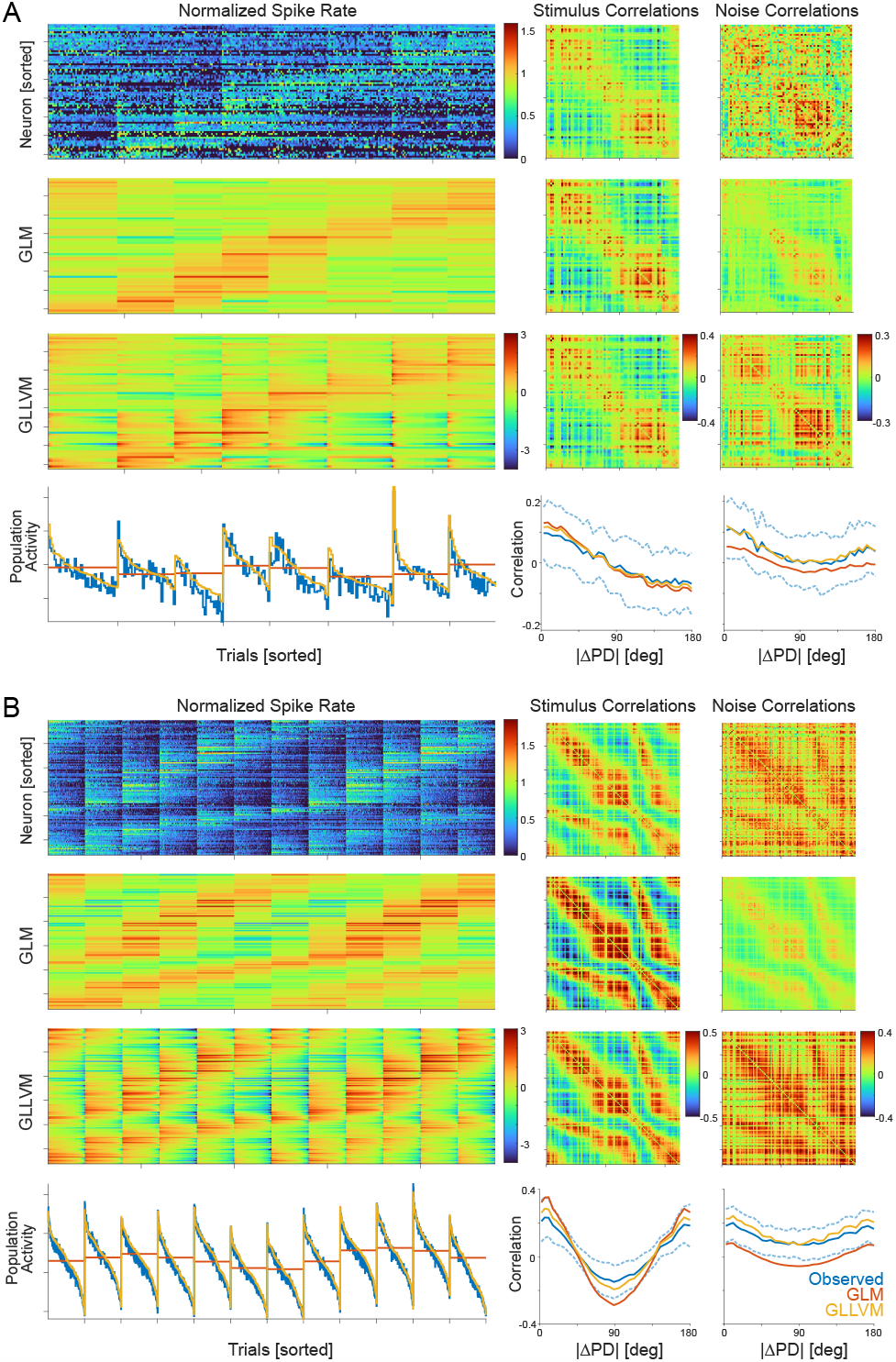
Encoding models for reach direction in M1 and grating direction from V1. A) Spike counts for all neurons recorded from M1 and trials for each of 8 directions of a center-out reaching task (top). Neurons are sorted by their preferred directions, and trials are sorted first by the target direction and then by the value of the latent state. The color scale is transformed 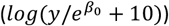 to highlight the differences across neurons and trials. Model fits for the GLM (Poisson observations) and GLLVM (1D latent, Poisson observations) are shown below, as well as the population activity. The observed and modeled stimulus and noise correlations are shown at right. B) Spike counts and model fits for neurons recorded from V1 responding to drifting full-field gratings in 12 directions (sorted as in A).

The dimensionality of the latent variable may have some impact on the encoding and decoding accuracy and on the calibration of Bayesian decoders. To characterize the potential effects of dimensionality we fit GLLVMs with 1 to 5 dimensional latent states for the M1, V1, and ABI datasets. We find that, in most cases, the GLLVMs with >1 dimensionality have similar error and coverage to the models with 1 dimension, with the exception of the Poisson GLLVM, which tends to have better coverage with more dimensions (Fig 6). In all cases the coverage of the NB models is better than that of the Poisson models.

**Figure 6:**
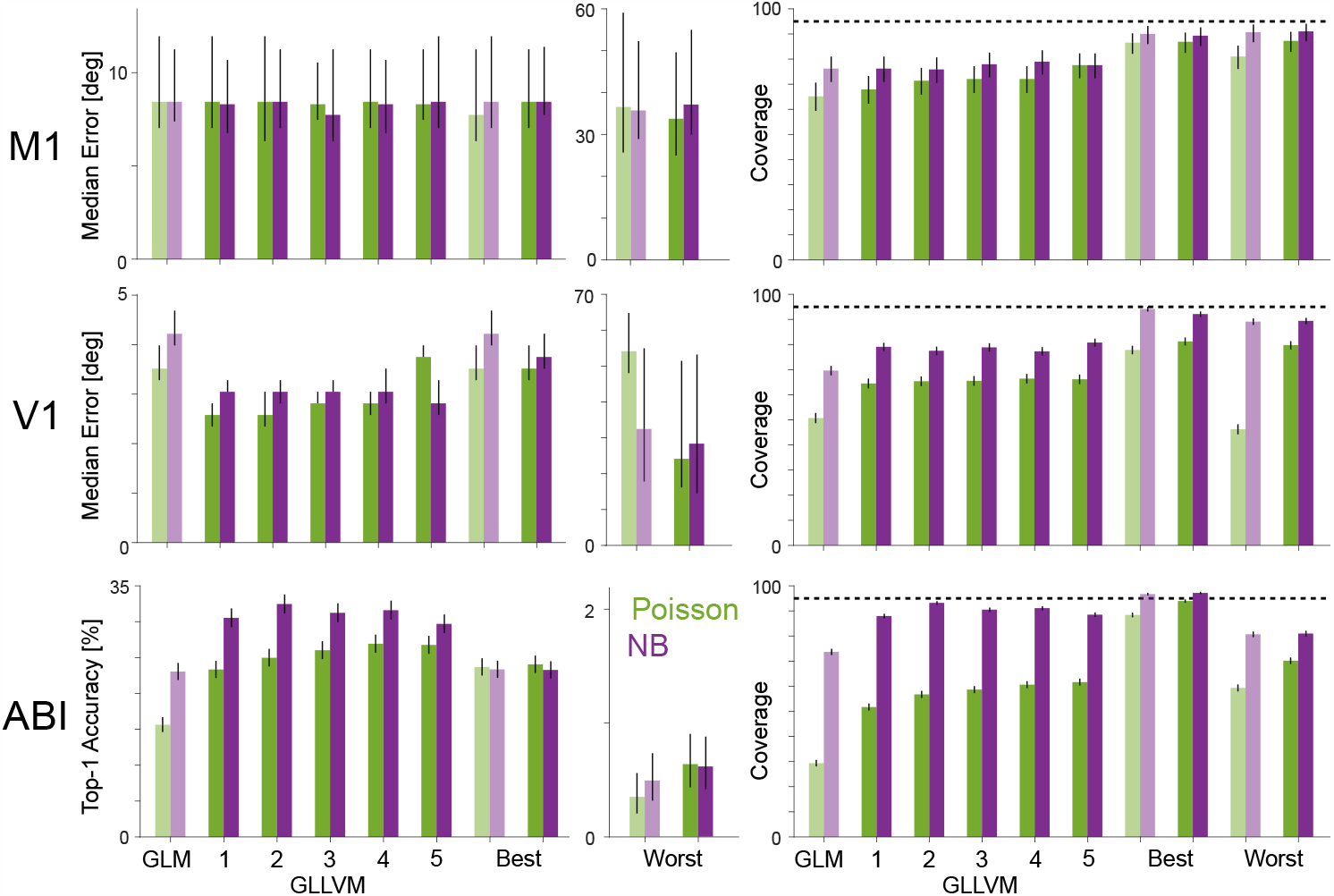
Increasing latent dimensionality does not fully correct over-confidence. Error/accuracy (left) and coverage at 95% credibility level (right) for GLLVMs with different latent dimensionality. GLM and GLLVM results reflect the full population of neurons for each experimental setting. For comparison, results with reduced populations of 20 neurons are included here for the GLM and one-dimensional GLLVM, selected using a greedy optimization to create the “best” and “worst” error/accuracy. Error bars denote 95% confidence intervals. Dashed lines denote nominated coverage of 95%. Light and dark colors for the best/worst greedy decoders denote results from the GLM and 1D GLLVM, respectively.

Since the size of the population appears to have an impact on coverage, we also examine how the composition of the population impacts accuracy and decoding. Here we use a greedy optimization (see Methods) to find the population of size N neurons that minimizes the error (M1 or V1) or maximizes the top-1 accuracy (ABI) of the Poisson GLM creating the greedy “best” subpopulation. And for comparison we also consider maximizing the error (M1 or V1) or minimizing the top-1 accuracy (ABI) of the Poisson GLM to create the greedy “worst” subpopulation. Like previous studies, we find that the full population is often unnecessary for accurate decoding – a greedy best subpopulation of N=20 often has error/accuracy comparable to the full population. Here we additionally show that these greedy best models often have better coverage than the models based on the full population (Fig 6). However, the population size is not the only factor determining coverage, since the greedy best and greedy worst populations have substantial differences in coverage despite both consisting of 20 neurons.

### Post-hoc correction for miscalibration

Since even decoders based on GLLVMs are over-confident, it may be useful to consider calibration as a distinct step in neural data analysis in situations where accurate uncertainty estimation is needed. One approach to correcting calibration errors is to simply inflate the posterior uncertainty post-hoc. That is, rather than decoding using *p*(*x*|*y*) use *q*(*x*|*y*) . Here we consider the transformation *q*(*x*|*y*) ∝ exp(*h* log *p*(*x*|*y*)) with *h* > 0. This transformation preserves the MAP estimate and the relative log-probabilities of all *x*, but *h* allows the uncertainty to be modified. Note that if *p*(⋅) is a normal distribution with standard deviation *σ, q*(⋅) is a normal distribution with standard deviation *σ*/√*h*, but this transformation can be used for general distributions.

For the over-confident examples above, we estimate a single constant *h* for each case (see Methods) and find that this transformation produces well-calibrated decoding distributions at all desired confidence levels (Fig 7A-C). The transformation does not change the decoding accuracy (based on MAP estimates) but allows for substantially more accurate uncertainty estimation. In the examples above, we showed that over-confidence depends on the encoding model and the number of neurons used in the decoder. The optimal value of *h*, thus, also depends on the model as well as the size and composition of the population with higher overconfidence needing greater correction (smaller *h*). We also note that, at least in some cases, underconfidence is possible (Fig 7D), but can be similarly corrected by *h* > 1.

**Figure 7:**
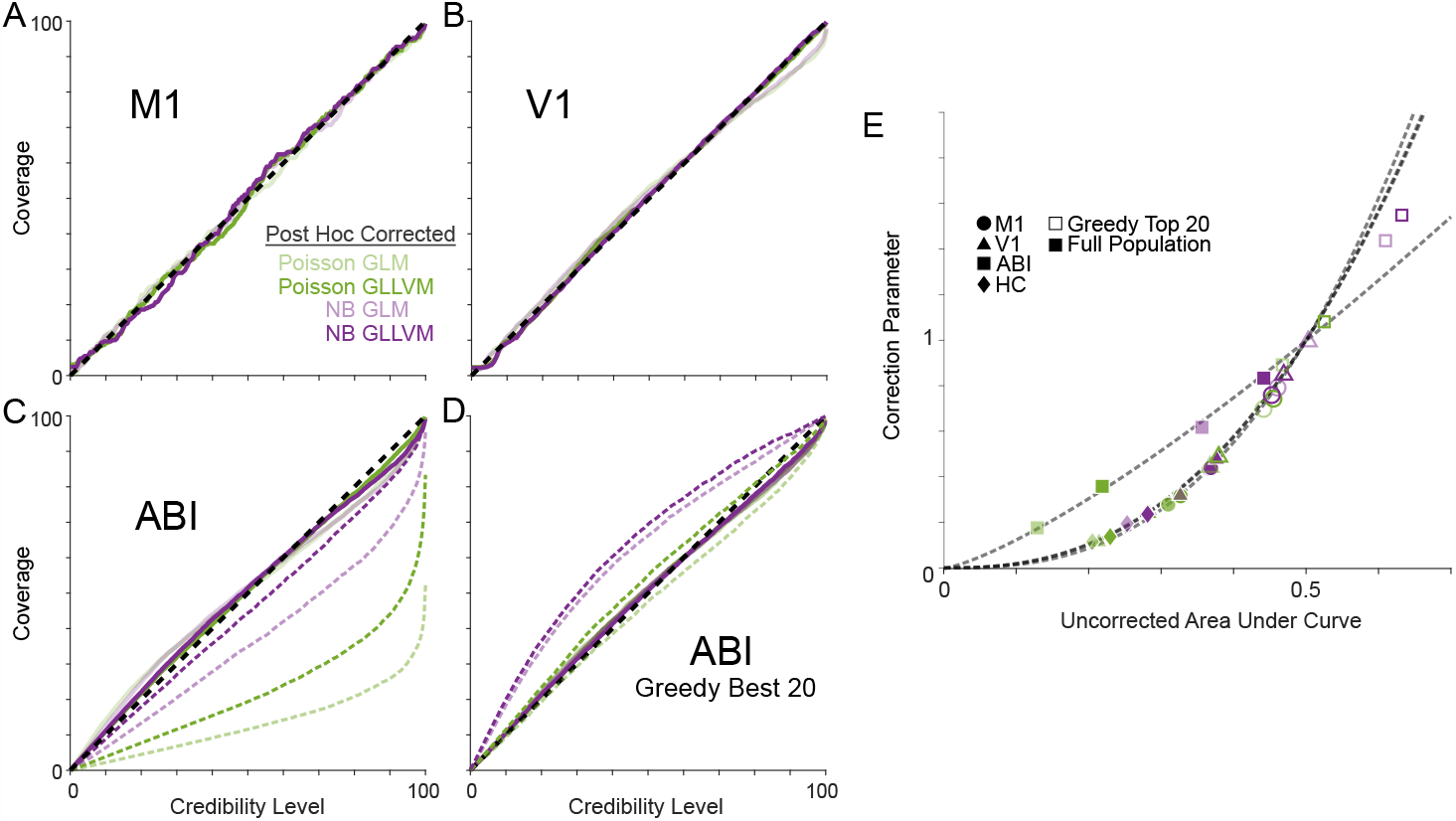
Post hoc corrected coverage. (A-C) results for full populations in each of the three experimental settings from Fig 3A-C. For each model and experiment there is a distinct correction parameter optimized to produce well calibrated results. D) Under-confidence is rare, but can occur, such as when decoding from the best 20 neurons (greedy selection) from the ABI dataset using NB models. Dashed lines in C and D decode the uncorrected results, while solid lines denote the post-hoc corrected results (dashed lines in C are repeated from 4C for reference). E) The optimal correction parameter as a function of original miscalibration. Dashed lines denote power law fits for each dataset.

Within a given experimental setting, there is a consistent relationship between the degree of over/under-confidence and the optimal correction parameter (here optimized by minimizing the mean squared error in the nominated coverage vs empirical coverage plots). Across models (GLM, GLLVM, Poisson, and NB) and populations (full population and greedy best), the correction parameters are well predicted by a power law, *h* = (2*p*)^K^, where *p* denotes the area under the curve for the uncorrected coverage and we find *a*d =2.7, 2.5, 1.3, 2.5 for M1, V1, ABI, and HC (see below), respectively (Fig 7E).

For some settings, rather than trial-by-trial decoding of spike counts, the goal is to decode a continuous, typically smoothly varying, external variable. To illustrate how general the problem of over-confidence in Bayesian decoders is, we consider continuous estimates of an animal’s position from hippocampal activity (Fig 8A). Here, rather than distinct trials with a controlled stimulus/behavior, a rat runs freely on a linear track. GLM and GLLVMs can still be used to decode the animal’s position. We fit encoding models based on place fields (direction-selective cubic B-spline bases with 10 equally spaced knots), and for the GLLVMs, we additionally include a one-dimensional latent variable. However, to more accurately decode the continuous behavior, we also add a process model that ensures that the position and latent state vary smoothly from one time to the next (see Methods).

**Figure 8:**
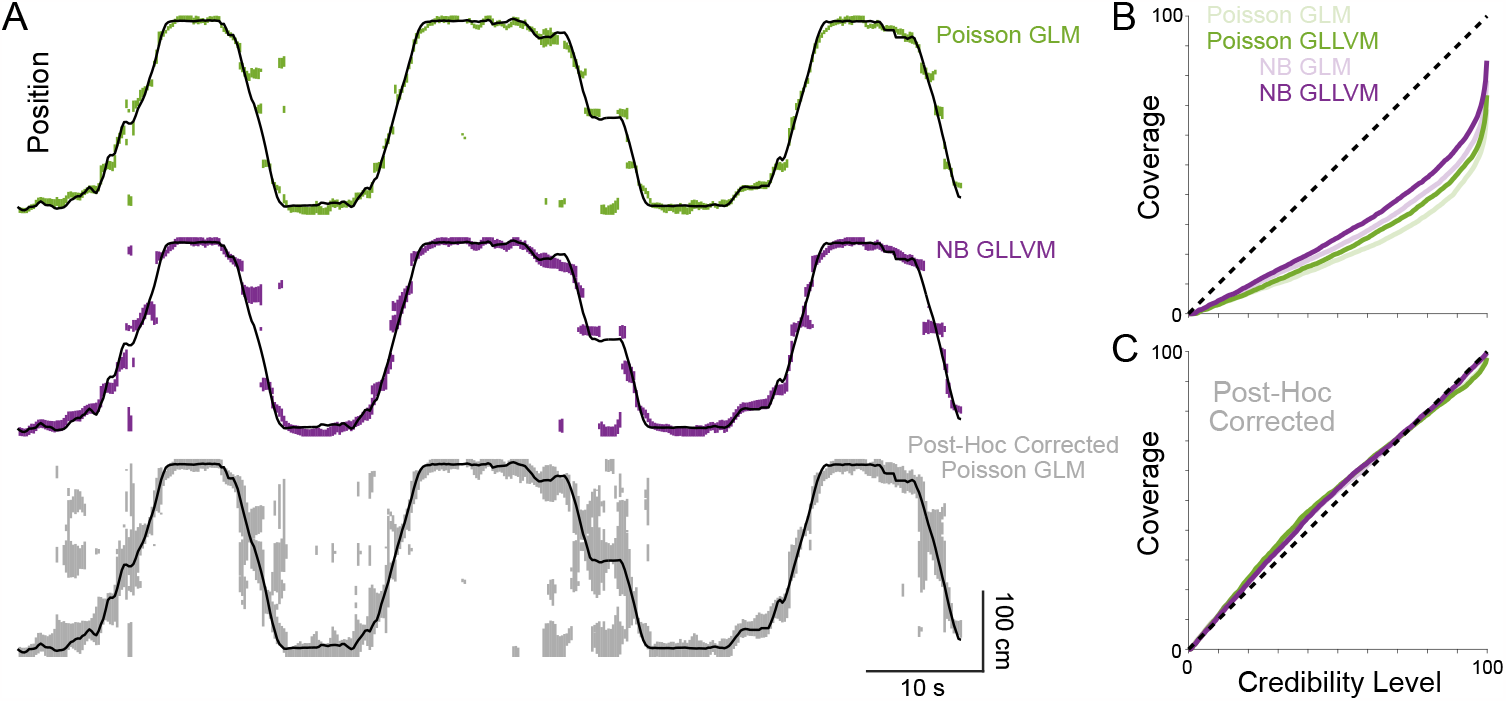
Continuous decoding and coverage for position in hippocampus (HC). A) The true position along the linear track (black line), along with 95% credible regions for three Bayesian decoders: 1) the traditional Poisson GLM, 2) a negative binomial GLM, and 3) the Poisson GLM after post-hoc correction. Note that, in some cases, the posterior (or post-hoc corrected distribution) is multimodal, resulting in multiple HPD regions. B) Empirical coverage as a function of the desired credibility level for the four Bayesian decoders. C) Empirical coverage after post-hoc correction.

As before, we assess the coverage of each model. Here we find that, decoding the time series of animal position, the Poisson GLM is the most overconfident and the NB-GLLVM is the most well-calibrated. The 95% credible regions for the posterior include the true position only 48% of the time for Poisson GLM, while the NB-GLLVM covers the true position 63% of the time (Fig 8B). All four models have better calibrated posteriors following post-hoc correction (Fig 8C). The coverage of 95% credible regions increases to 91% for the P-GLM and 94% for the NB-GLLVM, for example.

### Posterior uncertainty and task variables

From trial to trial there are substantial variations in both posterior uncertainty and accuracy. The exact relationship between error/uncertainty and accuracy depends somewhat on the decoder, since different models have different uncertainties. However, in the data examined above, we find that for all models error increases with increasing posterior uncertainty (M1 and V1) or accuracy decreases with increasing posterior uncertainty (ABI) (Fig 9). Fitting a linear model (in the log-log domain) for the post-hoc corrected Poisson GLM, M1 error increases 252% [187, 340] (95% CI) for each doubling of posterior (circular) standard deviation. For V1 with the post-hoc corrected Poisson GLM, error increases 160% [150, 169] for each doubling of the posterior (circular) standard deviation. Fitting a logistic model for ABI, accuracy decreases with OR=0.75 [0.68, 0.83] per bit of posterior entropy. These results are for the posteriors of the post-hoc corrected Poisson GLM, but all models show statistically significant dependencies between error/accuracy and uncertainty both with and without post-hoc correction.

**Figure 9:**
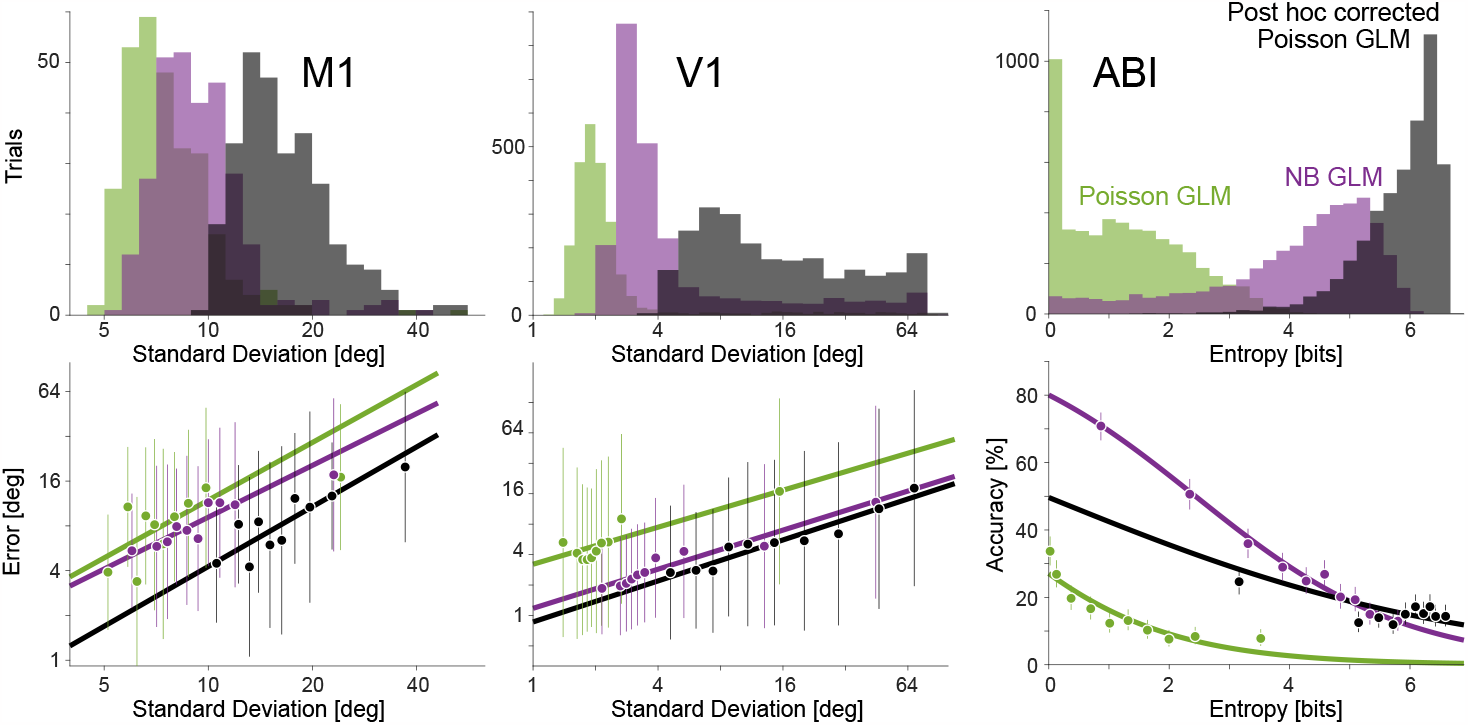
Uncertainty predicts accuracy. For reference, dots denote averages calculated in deciles. Error bars for M1 and V1 denote standard deviation. Error bars for ABI denote 95% confidence intervals. Lines for M1 and V1 denote linear, least-squares fit for single trials in the log domain. Curves for ABI denote logistic regression.

In experiments where a task variable is expected to influence behavioral/perceptual uncertainty, we may also expect Bayesian decoders to reflect differences in this uncertainty. Here, for instance, we examine V1 data from an additional experiment with static oriented grating stimuli, where the contrast of the stimulus was explicitly varied. Fitting separate (categorical) Poisson GLMs to the different time points (50ms window) and contrast conditions, we find that accuracy for decoding categorical stimulus orientation increases following stimulus onset and increases with increasing stimulus contrast (Fig 10A top). Accuracy for the high contrast trials is substantially higher than for low contrast trials (66% for high, 43% for low, z=7.4, p<10^-12^, two-sided test for difference of proportions, 200ms following stimulus onset). Additionally, posterior entropy decreases following stimulus onset, and is lowest for high contrast stimuli (Fig 10A middle). In this example, since the population is relatively small (18 units), the degree of over-confidence for the Poisson GLM (Fig 10A bottom) is not as extreme as the previous V1 population. Here, the post-hoc corrected posteriors for the Poisson GLM (corrected separately for each time point and contrast) show a similar pattern with high contrast trials having lower entropy than low contrast trials (1.3 bits for high, 2.1 bits for low, two-sided unpaired t-test t(955.4)=21.0, p<10^-12^, at 200ms following stimulus onset). As in Fig 9, we find that single trial accuracy is well predicted by the posterior uncertainty (Fig 10B). The relationship between entropy and accuracy is consistent across contrasts, and the logistic fits do not differ substantially for the different contrasts (OR=0.18/bit [0.12, 0.27] 95% CI for high contrast, OR=0.21/bit [0.14, 0.31] for low contrast).

**Figure 10:**
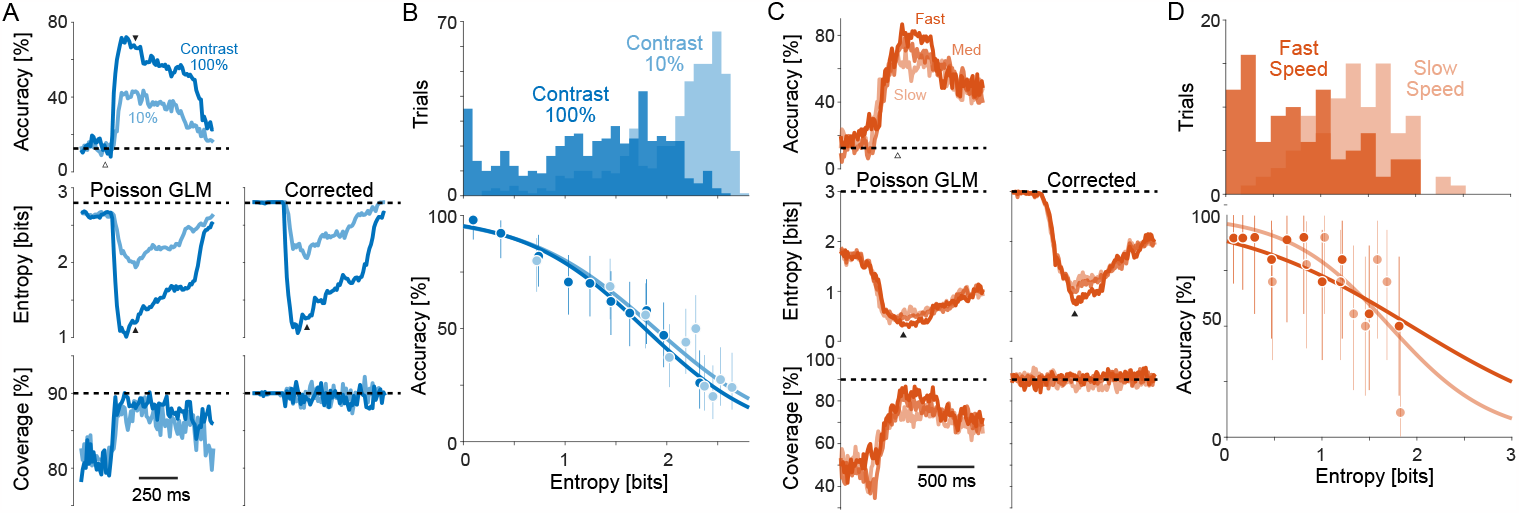
Accuracy, uncertainty, and coverage vary with stimulus contrast in V1 and with movement speed in M1. A) For static, oriented gratings, cross-validated decoding accuracy increases following stimulus onset (white triangle) but depends on stimulus contrast (top). Posterior entropy decreases, with lower entropy for higher contrast stimuli, and coverage (at 90% nominated) also varies. Dashed lines denote chance (top), maximum entropy (middle), and nominated coverage (bottom). B) At 200ms after stimulus onset (black triangles in A), we find that the (post hoc corrected) posterior entropy for the Poisson GLM varies with contrast. Dots denote averages in deciles, error bars denote 95% confidence intervals, and curves denote logistic regression fits. C, D) Analogous results for recordings from M1 during center-out reaching with maximum movement speed split by terciles. Cross-validated decoding accuracy increases shortly before movement onset (white triangle) but depends on reach speed (top). Posterior entropy decreases with lower entropy for higher speeds. Results in (D) are for 100ms after movement onset (black triangles in C).

We use a similar analysis to assess the impact of reach speed in M1. Just as stimulus contrast may impact uncertainty when decoding visual stimuli, movement features beyond reach direction may impact uncertainty when decoding behavior. Here we use the M1 data during center-out reaching examined above. We fit a single decoder for reach direction at each time point (50ms window), but assess accuracy, entropy, and coverage separately for different trials based on the peak movement speed. Splitting the trials into speed terciles (Fig 10C), we find that accuracy increases shortly before movement onset, and trials with the fastest reaches are decoded more accurately than those with slower reaches (80% for fast, 64% for slow, z=2.6, p=0.01, two-sided test for difference of proportions, 100ms following movement onset). Posterior entropy also decreases shortly before movement onset and is lowest for the fast reaches (Fig 10C middle). Here, as before, the Poisson GLM tends to be overconfident. The post-hoc corrected posteriors have substantially higher entropy, but show the same pattern where fast reaches have the lowest entropy (0.8 bits for fast, 1.4 bits for low, two-sided unpaired t-test t(184.5)=7.8, p<10^-12^, at 100ms following movement onset). The entropy on single trials again predicts single trial accuracy (Fig 10D), and the logistic fits do not differ substantially for the different speeds (OR=0.16/bit [0.06, 0.44] 95% CI for fast, OR=0.35/bit [0.13, 0.94] for slow).

## Discussion

Using data from a range of brain regions and experimental settings, we have shown how Bayesian decoders of neural spiking activity are often miscalibrated. In particular, the posterior estimates tend to be overconfident. Overconfidence increases with increasing numbers of neurons, is reduced by using negative binomial observation models (compared to Poisson), and is reduced by modeling latent variables. However, since even the best calibrated models tested here are not well calibrated, we introduce a post-hoc correction and show how it can be used, in multiple settings, to recalibrate uncertainty estimates. Finally, we present results illustrating how the posterior uncertainty of Bayesian decoders can vary substantially from trial-to-trial. Single trial posterior uncertainty predicts single trial accuracy and may be useful for understanding variation in perceptual or behavioral confidence due to task variables such as stimulus contrast or movement speed.

Similar to previous work (Macke et al., 2011), we show here how latent variables (GLLVMs) can better account for noise correlations and shared variability in the simultaneously recorded neurons. Correlations are known to play an important role in population coding, generally (von der Malsburg, 1994; Nirenberg, 2003), and failing to accurately account for these dependencies can lead to decoding errors (Ruda et al., 2020). Latent variable models represent one approach to describing shared variability, and previous work has shown how these models can improve encoding and decoding accuracy (Santhanam et al., 2009; Chase et al., 2010; Lawhern et al., 2010). Here we additionally show how latent variable models increase the uncertainty of Bayesian decoders and improve their calibration.

Bayesian decoders have advantages over other decoding methods in that they provide probabilistic predictions and can flexibly incorporate prior assumptions, such as sparseness and smoothness. However, many non-Bayesian decoders exist, including vector decoders (Georgopoulos et al., 1986; Salinas and Abbott, 1994), nearest-neighbor methods, support vector machines, and artificial neural networks (Quiroga and Panzeri, 2009). Although, well-tuned Bayesian methods can often out-perform non-Bayesian approaches (e.g. Zhang et al., 1998). Machine learning and recent deep learning approaches to decoding have been shown to be more accurate than simple Bayesian models in many settings (Pandarinath et al., 2018; Glaser et al., 2020b; Livezey and Glaser, 2021). Since calculating the full posterior distribution can be computationally expensive, these methods can also be substantially faster for situations where predictions are time-sensitive. Almost all work with non-Bayesian decoders of neural activity focuses on the accuracy of point predictions. However, miscalibration is a known problem in work on artificial neural networks (Guo et al., 2017) and recent work on Bayesian neural networks and conformal prediction (Shafer and Vovk, 2008) could potentially be used to create and calibrate uncertainty estimates for these models.

Accurate uncertainty estimates may potentially be useful for robust control of brain machine interfaces (BMIs). For instance, although many BMIs directly control effectors, such as a cursor position (decoding movement) or a desired word (decoding speech), based on point predictions (Nicolelis, 2003), it may be beneficial to distinguish between predictions based on their confidence level. Here, we find substantial variation in uncertainty for trial-by-trial offline decoding, and we also illustrate how contrast (in V1) and speed (in M1) might impact decoding uncertainty. These results are limited by the fact that we do not explicitly include contrast or speed in the encoding model (Moran and Schwartz, 1999) or decode these variables directly (Inoue et al., 2018), but they suggest how uncertainty may be a separate and worthwhile consideration for decoding problems. Additionally, our results suggest that recalibration could be necessary to avoid overconfidence in BMIs that make use of posterior uncertainty during control.

The uncertainty estimates from Bayesian decoders of neural activity may also be useful for studying behavioral and perceptual uncertainty. Normative models of population coding (Ma et al., 2006) and broader descriptions of uncertainty in the brain (Knill and Pouget, 2004) often directly relate neural activity to probabilistic descriptions of the external world. Although several features of neural activity have been proposed as indicators of behavioral/perceptual uncertainty (Vilares and Kording, 2011), the posteriors from Bayesian decoders represent a principled framework for translating noisy, high-dimensional data into a single probabilistic description (Zemel et al., 1998; Dehaene et al., 2021; Kriegeskorte and Wei, 2021). The impacts of tuning curve shapes (e.g. Pouget et al., 1999; Zhang and Sejnowski, 1999) and correlations between neurons (Averbeck et al., 2006; Lin et al., 2015; Kohn et al., 2016) on the uncertainty of population coding have been well studied, and here we add to this work by demonstrating how different encoding models (GLM vs GLLVM and Poisson vs negative binomial) have systematically different degrees of overconfidence in experimental recordings across many settings.

Since even the best Bayesian models (negative binomial latent variable models up to five dimensions) are overconfident, recalibration appears to be necessary to ensure that the uncertainty of Bayesian decoders matches the distribution of errors. On one hand, this may suggest that there is additional mismatch between the GLLVM and the data generating process. It may be that low-dimensional latent variable models only partially capture noise correlations (Stevenson et al., 2012), that there is unmodeled nonstationarity in the tuning curves (Cortes et al., 2012; Rule et al., 2019), that responses are underdispersed (DeWeese et al., 2003; Stevenson, 2016), or some combination of these factors. On the other hand, humans and other animals are often over-or underconfident during perceptual and cognitive judgements (Baranski and Petrusic, 1994; Kepecs and Mainen, 2012; Mamassian, 2016). It is possible that the original (miscalibrated) uncertainty estimates better predict psychophysical uncertainty or metacognitive reports of confidence, even if recalibrated uncertainty estimates better predict the distribution of external variables.

Finally, it is important to note that when Bayesian models are recalibrated post-hoc they are no longer following a coherent Bayesian framework (Dawid, 1982). From a practical standpoint, such as when developing BMIs, model calibration may be more important than model coherence. However, additional work is needed to better understand the alignment of perceptual/behavioral uncertainty and decoder posterior uncertainty (Panzeri et al., 2017). Models with more accurate descriptions of single neuron variability (Gao et al., 2015; Ghanbari et al., 2019), with nonstationarity (Shanechi et al., 2016; Wei and Stevenson, 2023), additional stimulus/movement nonlinearities (Schwartz and Simoncelli, 2001), state-dependence (Panzeri et al., 2016), and with more complex latent structure (Glaser et al., 2020a; Williams et al., 2020; Sokoloski et al., 2021; Williams and Linderman, 2021) may all show better coverage while maintaining coherence. Our results here indicate that Bayesian decoders of spiking activity are not necessarily well calibrated by default.

## Acknowledgments

This material is based upon work supported by the National Science Foundation under Grant 1931249 and Grant 1848451.

